# High precision registration between zebrafish brain atlases using symmetric diffeomorphic normalization

**DOI:** 10.1101/081000

**Authors:** Gregory D. Marquart, Kathryn M. Tabor, Eric J. Horstick, Mary Brown, Harold A. Burgess

## Abstract

Atlases provide a framework for information from diverse sources to be spatially mapped and integrated into a common reference space. In particular, brain atlases allow regional annotation of gene expression, cell morphology, connectivity and activity. In larval zebrafish, advances in genetics, imaging and computational methods have enabled the collection of large datasets providing such information on a whole-brain scale. However, datasets from different sources may not be aligned to the same spatial coordinate system, because technical considerations may necessitate use of different reference templates. Two recent brain atlases for larval zebrafish exemplify this problem. The Z-Brain atlas contains information on gene expression, neural activity and neuroanatomical segmentation acquired using immunohistochemical staining of fixed tissue. In contrast, the Zebrafish Brain Browser (ZBB) atlas was constructed from live scans of fluorescent reporter genes in transgenic larvae. Although different reference brains were used, the two atlases included several transgene patterns in common that provided potential 'bridges' for transforming each into the other’s coordinate space. We therefore tested multiple bridging channels and registration algorithms. The symmetric diffeomorphic normalization (SyN) algorithm in ANTs improved the precision of live brain registration while better preserving cell morphology than the previously used B-spline elastic registration algorithm. SyN could also be calibrated to correct for tissue distortion introduced during fixation and permeabilization. Finally, multi-reference channel optimization provided a transformation matrix that enabled Z-Brain and ZBB to be co-aligned with acceptable precision and minimal perturbation of cell and tissue morphology. This study demonstrates the feasibility of integrating whole brain datasets, despite disparate acquisition protocols and reference templates, when sufficient information is present for bridging.

**Anatomical abbreviations:**

ac: anterior commissure
DT: Thalamus
GT: Griseum tectale
Ha: Habenula
Hc: Hypothalamus caudal zone
Hi: Hypothalamus intermediate zone
MO: Medulla oblongata
NXm: Vagus motor neurons
OB: Olfactory bulb
OE: Olfactory epithelium
IO: Inferior olive
LC: Locus coeruleus
MN: Mauthner neuron
MO: Medulla oblongata
Pal: Pallium
pc: posterior commissure
Pr: Pretectum
SR: Superior raphe
Teg: Tegmentum
TeOn: Optic tectum neuropil
TG: Trigeminal ganglion
TL: Torus longitudinalis

## Introduction

Larval stage zebrafish are an increasingly popular model for neurobiological studies. With a brain that contains an estimated 10^5^ neurons, larvae are similar in complexity to adult *Drosophila*, another established neuroscience model. In both systems, researchers can deploy a wide range of genetic and transgenic tools in efforts to decode patterns of neural structure and connectivity. In larval zebrafish, optical transparency and constrained physical dimensions (fitting within an imaging volume of 1000 × 600 × 350 μm) allow the entire brain to be rapidly scanned at cellular resolution using diffraction-limited microscopy. In principle, this enables researchers to systematically analyze effects of manipulations on a brain-wide level. However, such efforts have been hampered by the absence of a comprehensive digital atlas that would provide researchers with a unified framework in which to aggregate data from different experiments and gain deeper insights from correlations between neuronal cell identity, connectivity, gene expression and function within the brain. Additionally, digital atlases may better define structural boundaries within the brain that are difficult to clearly identify within individual instances, allowing for a more rigorous mapping of neuroanatomical regions onto experimental data.

This longstanding problem in zebrafish neuroscience has recently been addressed by the construction of digital atlases using three-dimensional image registration techniques: the Virtual Brain Explorer for Zebrafish (ViBE-Z), Z-Brain and the Zebrafish Brain Browser (ZBB) (Marquart et al., 2015; Randlett et al., 2015; Ronneberger et al., 2012). In these atlases, information on gene expression, structure (neuronal cell bodies, glia, vasculature, ventricles, neuropil or axon tracts) and measures of activity (calcium or secondary messenger activity) are consolidated within a common spatial framework. By using widely-available transgenic lines or immunohistochemical stains as reference templates for brain alignment, each of these atlases provides other researchers the opportunity to register their own datasets into these digital spaces and take advantage of the information contained within.

ViBE-Z was the first comprehensive three-dimensional digital brain atlas in zebrafish that used a nuclear stain for the alignment of 85 high resolution scans comprising 17 immunohistochemical patterns at 2-4 days post-fertilization (dpf) (Rath et al., 2012; Ronneberger et al., 2012). In ViBE-Z, custom algorithms were developed to correct for variations in fluorescent intensity with scan depth, and a landmark approach taken to perform accurate image registration and segmentation into 32 brain regions.

In contrast, two more recent approaches (Z-Brain and ZBB) have generated brain atlases at 6 dpf through non-linear B-spline registration using the freely available Computational Morphometry Toolkit (CMTK) (Portugues et al., 2014; Rohlfing and Maurer, 2003). Z-Brain includes 29 immunohistochemical patterns from 899 scans which form the basis for expert manual segmentation of the brain into 294 anatomical regions. These partitions facilitate the analysis of phospho-ERK expression for mapping neural activity (Randlett et al., 2015). In Z-Brain, each expression pattern was co-scanned with tERK immunoreactivity, and registered to a single tERK-stained reference brain. For ZBB, we live-imaged 354 brains from 109 transgenic lines and manually annotated the expression found in each (Marquart et al., 2015). In place of tERK, a single *vglut2a:dsRed* transgenic brain was used as the reference in ZBB with every transgenic line crossed to this transgenic line to provide a channel for registration. ZBB enables researchers to select a transgenic line labeling a selected set of neurons for monitoring and manipulating circuit function.

While Z-Brain and ZBB are powerful datasets on their own, we saw an opportunity to bridge the two atlases as they are both based on confocal scans of 6 dpf larvae. This would bring to Z-Brain a large number of additional transgenic lines and to ZBB, the expert manual segmentation of Z-Brain. Several similarities between Z-Brain and ZBB suggested that merging the atlases would be possible. First as zebrafish rearing conditions are standardized across laboratories and fish were imaged at the same time post-fertilization, Z-Brain and ZBB likely reflect the same developmental timepoint. Second, images in both atlases were acquired at similar resolution (0.8 × 0.8 × 2 μm for Z-Brain; 1 × 1 × 1 or 1 × 1 × 2 μm for ZBB) and orientation (dorsal to ventral horizontal scans). Third, despite using distinct templates (tERK for Z-Brain and *vglut2a* for ZBB), Z-Brain and ZBB share several transgenic markers in common, which provided the possibility of bridging the datasets by using these shared patterns as references for a secondary registration step.

One of the strengths of larval zebrafish is the ability to rapidly image the brain at cellular resolution and visualize brain-wide neuronal morphology, providing valuable information on cell type and potential connectivity. Z-Brain and ZBB both illustrate the feasibility of performing whole-brain registration with precision sufficient to ensure that the ’same' neurons from different fish are aligned to within a cell diameter (~10 μm). However, a challenge for brain registration in zebrafish is to minimize local distortions, so that cellular morphology is preserved while still allowing sufficient deformation to overcome biological variability between individual brains or malformations due to tissue processing.

Here we describe a method we used to co-register ZBB and Z-Brain, bridging the two existing 6 dpf larval zebrafish brain atlases. By using the diffeomorphic algorithm SyN in the software package Advanced Normalization Tools (ANTs) (Avants et al., 2008, 2011), we were able to overcome differences in tissue shape due to fixation, optimize the trade-off between preservation of cell morphology and global alignment, and provide precise registration in all brain regions. Additionally, ANTs provided superior image registration for live-scanned larvae, enabling us to improve the precision of registration and neuron morphology within our original ZBB atlas, allowing us to compile a new version with improved fidelity (ZBB_1.2_).

## Methods

### Zebrafish lines

In order to provide additional options for bridging ZBB and Z-Brain, we scanned two transgenic lines that were not in the original ZBB release: *Et(gata2a:EGFP)zf81* (*vmat2:GFP*) and *Tg(isl1:GFP)rw*0 (*isl1:GFP*) (Higashijima et al., 2000; Wen et al., 2008). Imaging and registration was performed as previously described (Marquart et al., 2015). Other lines referenced in this study are *Tg(slc6a3:EGFP)ot80* (*DAT:GFP*) (Xi et al., 2011), *Tg(-3.2fev:EGFP)ne0214* (*Pet1:GFP*) (Lillesaar et al., 2009), *y264Et* (Tabor et al., 2014), *s1181Et* (Baier and Scott, 2009), *Tg(gad1b:GFP)nns25* (*gad1b:GFP*) (Satou et al., 2013), *Tg(slc6a5:GFP)cf3* (*glyT2:GFP*) (McLean et al., 2007), *Tg(-17.6isl2b:GFP)zc7* (*isl2b:GFP*) (Pittman et al., 2008), *Tg(-3.4tph2:Gal4ff)y228* (*tph2:Gal4*) (Yokogawa et al., 2012), *TgBAC(slc17a6b:lox-DsRed-lox-GFP)nns14* (*vglut2a:dsRed*) (Satou et al., 2012), *Tg(slc17a6:EGFP)zf139* (Bae et al., 2009), *Tg(elavl3:CaMPARI(W391F+V3987L))jf9* (Fosque et al., 2015), *Tg(phox2b:GFP)w37* (Nechiporuk et al., 2007), *J1229aGt* (Burgess et al., 2009) and several Gal4 enhancer traps from ZBB: *y304Et, y332Et, y341Et, y351Et* and *y393Et* (Marquart et al., 2015).

### Immunohistochemistry

Immunolabeling was as described (Randlett et al., 2015) with the following adaptations. Larvae were fixed overnight at 4°C in PBS with 4% paraformaldehyde and 0.25% Triton X-100. Samples were then washed in PBS containing 0.1% Triton X-100 (PBT) for 5 min, 3 times. For antigen retrieval, samples were incubated in 150 mM Tris-HCl ph 9.0 for 5 min, followed by 15 min at 70°C and washed in PBT for 5 min, 2 times (Inoue and Wittbrodt, 2011). Critically, samples were then permeabilized on ice in fresh 0.05% trypsin-EDTA for no more than 5 minutes. If pigmented, samples were incubated in PBT with 1.5% H_2_O_2_ and 50 mM KOH for 15 min, rinsed 2 times in PBT and washed again for 10 min. Samples were then blocked in PBT containing 5% normal goat serum (NGS) and 0.2% bovine serum albumin (BSA) for 1 hr before incubation at 4°C with tERK antibodies (Cell Signaling, 4696) diluted 1:500 in PBT with 5% NGS and 0.2% BSA for a minimum of 6 hr. Samples were then washed with PBT for 30 min, 4 times, before incubation at 4°C for a minimum of 2 hr with fluorescent secondary antibodies (Alexa Fluor 488 or 548) diluted 1:1000 in PBT with 5% NGS and 0.2% BSA. Samples were finally rinsed for 30 min, 4 times, prior to imaging.

### Software

Registrations were performed using CMTK version 3.2.3 and ANTs version 2.1.0 running on the National Institute of Health’s Biowulf Linux computing cluster. Registrations were parallelized using Slurm-based bash scripts available upon request. For CMTK, previously optimized registration parameters that minimize computation time while maximizing precision were used (registrationx --dofs 12 --min-stepsize 1 and warpx --fast --grid-spacing 100 --smoothness-constraint-weight 1e-1 --grid-refine 2 --min-stepsize 0.25 --adaptive-fix-thresh 0.25). For ANTs registrations, the parameters used are cited in the relevant text and figures with optimized parameters listed in Table 1. Image volumes were rendered within the Zebrafish Brain Browser (ZBB), NIH Image (Schneider et al., 2012) or code written in IDL (www.harrisgeospatial.com). For the conversion to/from NIfTi format required for ANTs, we used the ImageJ plugin nifti_io.jar written by Guy Williams.

**Table 1.**
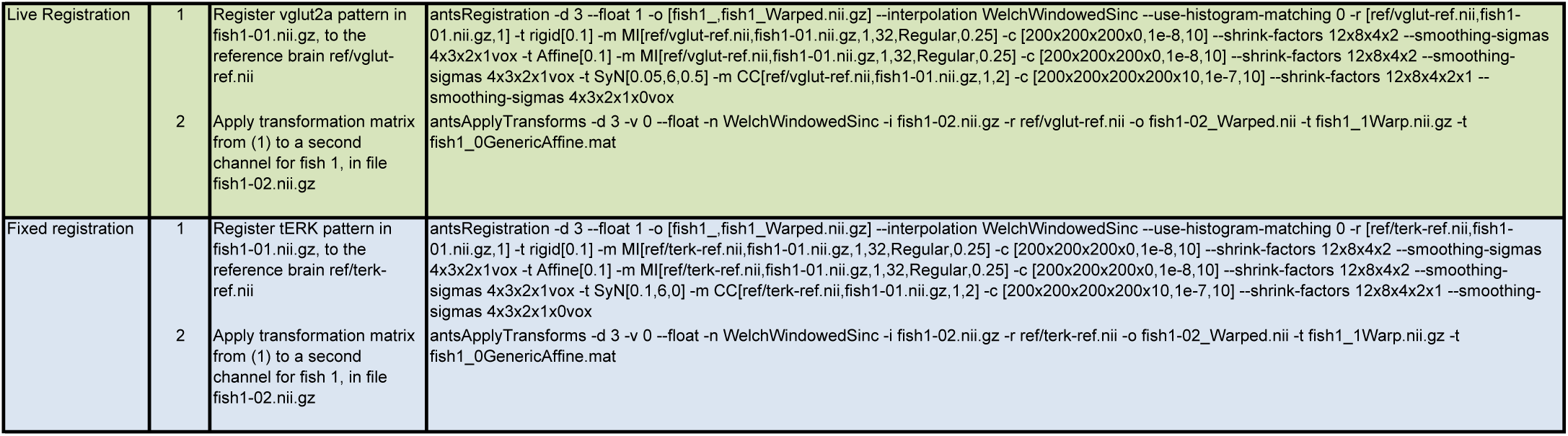
ANTs command parameters for image registration

### Statistics

Cross correlation between registered image sets was performed using the c_correlate function within IDL version 7.0. Correlations were run within small sub-regions of the registered image volumes. In Fig. 1,3 & 4, 50 μm side cube sub-regions were manually defined by selecting volumes containing high contrast boundaries. For cross correlations between individual brains scanned for each transgenic line in ZBB (Fig 2a,b), 40 μm side cubes were drawn around the three computationally identified brightest sub-regions within the expression pattern, with cross-correlation then calculated between all pairs of brains. The mean of all cross-correlations was used to estimate registration precision.

**Figure 1.**
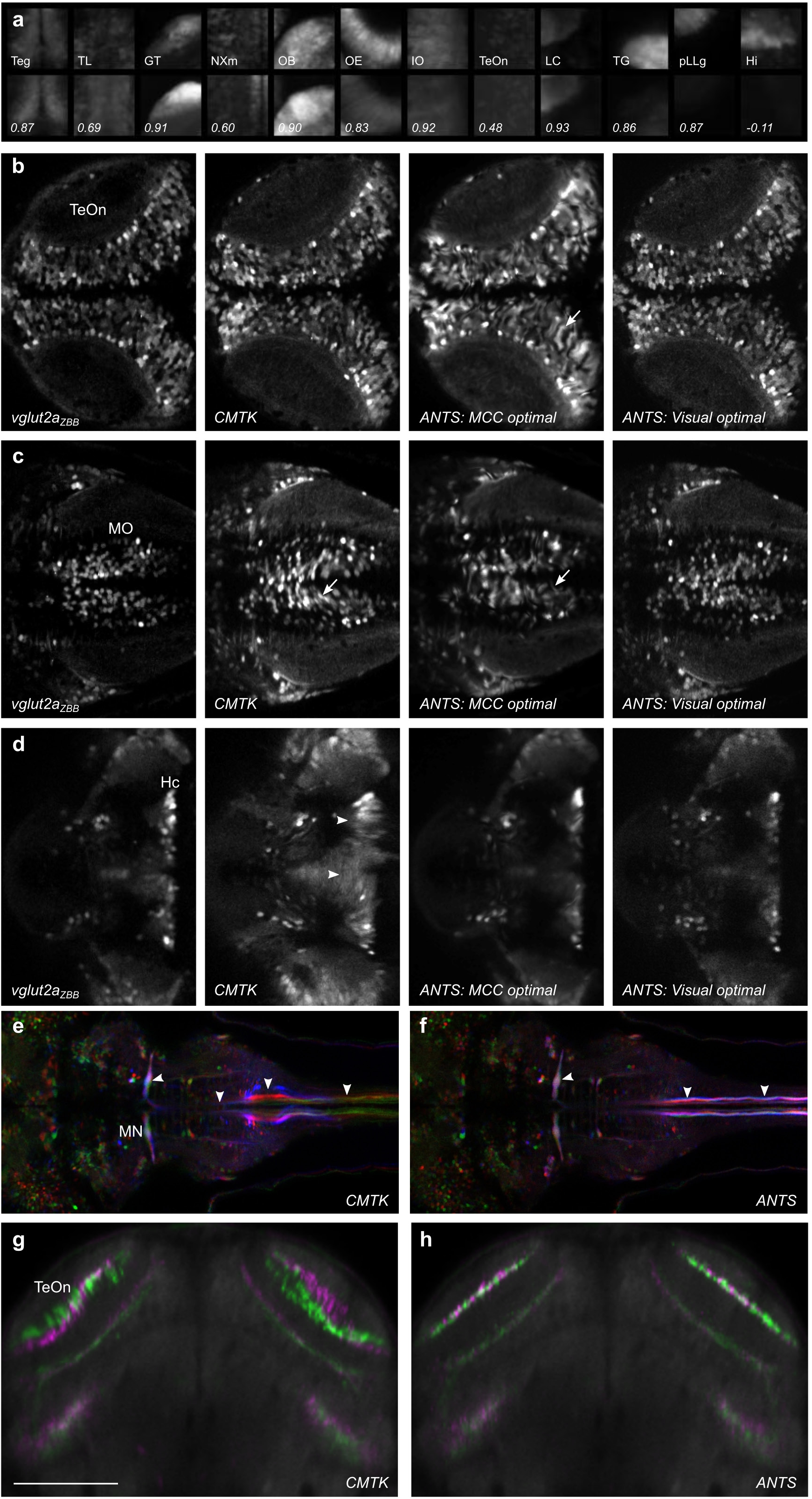
Optimization of parameters for registration of live brain scans using ANTs. (**a**) Dorsal maximum projections through the twelve 50 × 50 × 50 μm cubes used to calculate the mean cross correlation (MCC) for *vglut2a* expression patterns. Top row shows projections for the reference image, *vglut2a*_ZBB_, and bottom row shows projections for a representative *vglut2a:dsRed* brain that was registered to the reference brain using CMTK. Correlation coefficients are indicated in the bottom row. For this example, the MCC is the mean of the indicated values, 0.73. **(b-d)** Comparison of a single plane in *vglut2a*_ZBB_, and of the representative *vglut2a:dsRed* brain after registration using CMTK, ANTs with parameters that produced the largest mean cross correlation score (0.85; MCC optimal), and ANTs with parameters where visual inspection showed cell morphology was best preserved (Visual optimal. MCC was 0.81). Slices are through the optic tectum (**b**), medulla oblongata (**c**) and hypothalamus (**d**). Distortion artifacts introduced by CMTK in the hypothalamus (arrowhead) as well as poor cell morphology with CMTK and ANTs-MCC-optimal (arrow) are indicated. (**e,f**) Comparison of a single horizontal plane in *J1229aGt* showing expression of GFP in the Mauthner cell and axon (arrowheads) for three individual larvae (pseudo-colored red, green and blue). Registration was performed with CMTK (**e**) or ANTs (**f**). (**g,h**) Single coronal plane through the optic tectum in two separate average brain images (colored green and magenta) for *y393Et*. For each brain image, we independently scanned three individual brains and registered them using CMTK (**g**) or ANTs (**h**). Scale bar 100 µm.

**Figure 2.**
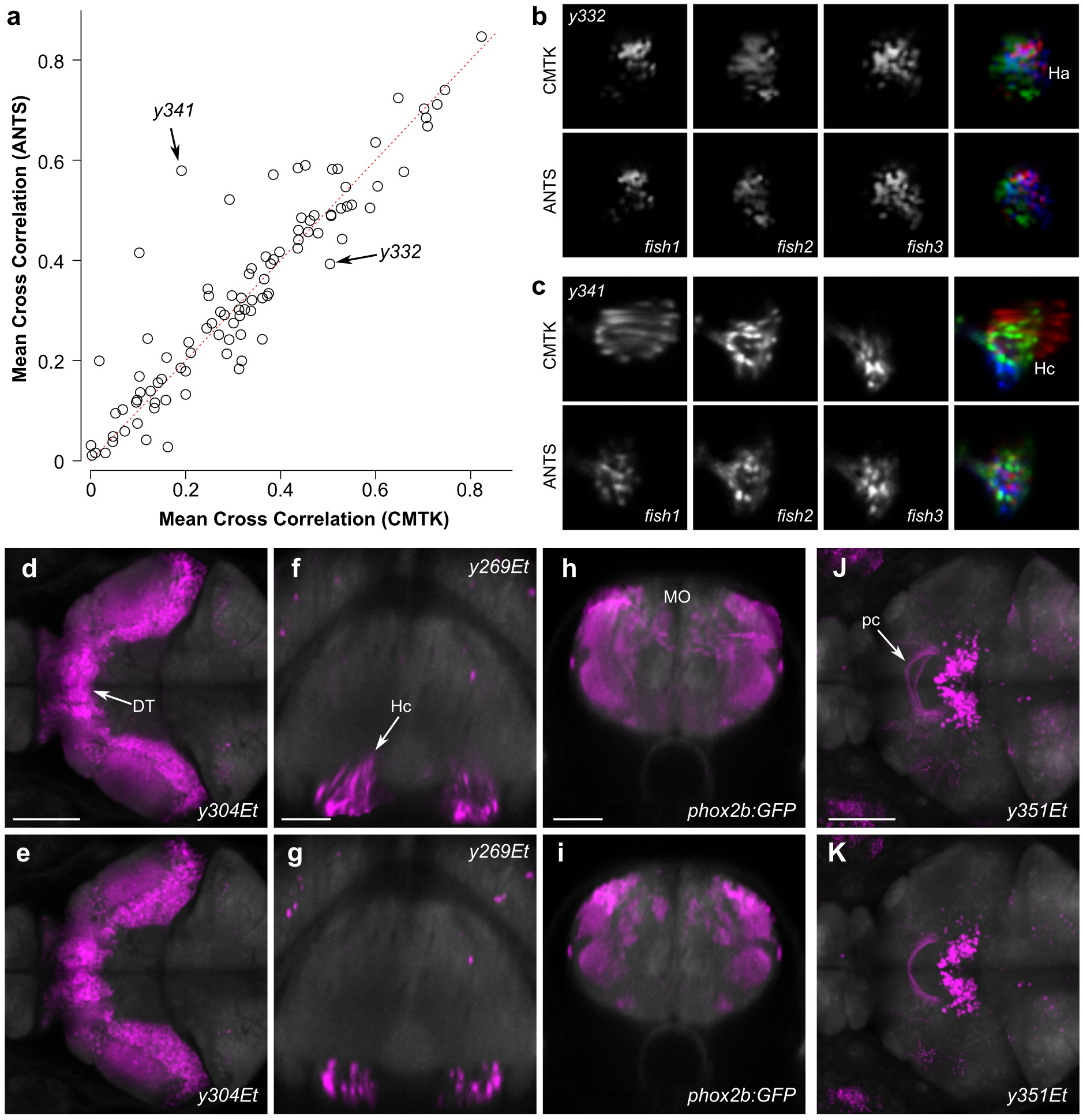
Improved precision of transgene representations in ZBB_1.2_. (a) Mean of cross-correlation values derived from all pairwise comparisons of individual brains for each transgenic line in ZBB, after registration with CMTK and ANTs. Dotted line indicates 1:1 ratio. (b) Horizontal slice through the right habenula in *y332Et*, showing three individual brain scans after registration with CMTK (top row), and the same slices pseudo-colored (red, green blue) and superimposed. Bottom row shows the equivalent after registration using ANTs. (c) Horizontal slice through the caudal hypothalamus of three individual *y341Et* larvae as well as their pseudo-colored superimposition following registration with CMTK (top row) or ANTs (bottom row). (**d,e**) Horizontal slice through the thalamus showing the averaged representation of enhancer trap line *y304Et*, where individual brains were registered with CMTK for ZBB (**d**), or by ANTs for ZBB_1.2_ (**e**). Arrow indicates neurons that are artificially elongated across the midline. Scale bar 100 µm. (**f,g**) Coronal slice through the caudal hypothalamus showing the average enhancer trap line *y269Et* brain with CMTK (**d**) and ANTs (**e**). Scale bar 50 µm. (**h,i**) Coronal slice through the medulla oblongata showing the average *phox2b:GFP* brain with CMTK (**f**) and ANTs (**g**). Scale bar 50 µm. (**j,k**) Horizontal projection through the posterior commissure (arrow) for the average *y351Et* brain obtained with CMTK (**j**) or ANTs (**k**). Scale bar 100 µm.

**Figure 3.**
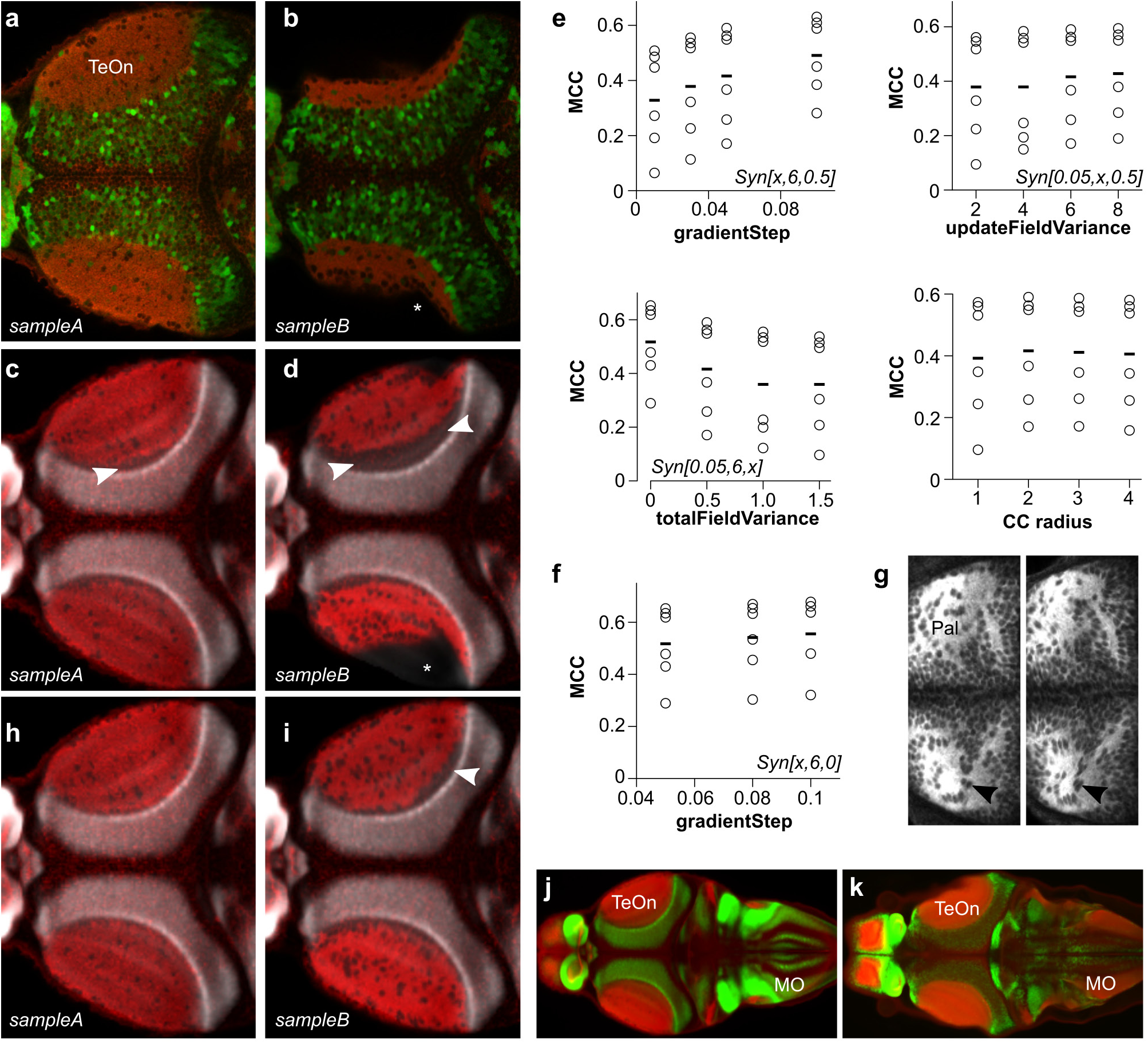
Optimization of ANTs registration parameters for fixed tissue. (**a,b**) Horizontal section through the optic tectum after immunostaining for tERK (red) and DsRed in *vglut2a:DsRed* (green), using diluted (**a, sample A**) or fresh trypsin (**b, sample B**). Asterisk indicates missing area of tectal neuropil due to permeabilization artifact. (**c,d**) Registration, using the *vglut2a:DsRed* expression pattern, of the tERK immunostain (red) in same brains as in (**a,b**) to tERK_ZBB_ using the parameters previously optimized for live registration. White shows the ZBB_1.2_ *vglut2a:dsRed* pattern. Arrowheads highlight regions where tERK in the optic tectum neuropil fails to closely abut the cellular layer. (e) Mean cross-correlation values for the tERK expression pattern after registration of 6 brains to tERK_ZBB_, varying each of the parameters for the ANTs SyN transform, starting with the parameters that gave the best registration for live *vglut2a:dsRed* based registration (Syn[0.05,6,0.5]). Bottom right: MCCs after varying the radius of the cross-correlation metric used during registration. (f) MCCs for tERK in the same brains as in (**e**), after combining the two best parameter sets from (**e**) (SyN[0.1,6,0.5] and Syn[0.05,6,0]) to assess further improvement in registration precision. (g) Horizontal section for comparison of tERK stain revealing cell morphology in the pallium after registration with optimal parameters for live *vglut2a* registration (left), and optimal parameters for registering fixed and stained tissue (right). (**h,i**) Same brains as in (**c,d**), but after registration to tERK_ZBB_ using the parameters optimized for fixed tissue registration. (**j,k**) Horizontal section through the optic tectum showing tERK expression (red) and *vglut2a:dsRed* expression (green) in ZBB_1.2_ (**j**) and Z-Brain (**k**). Matching slices within the optic tectum were selected; because the rotation around the y-axis is slightly different, sections are different within the medulla.

**Figure 4.**
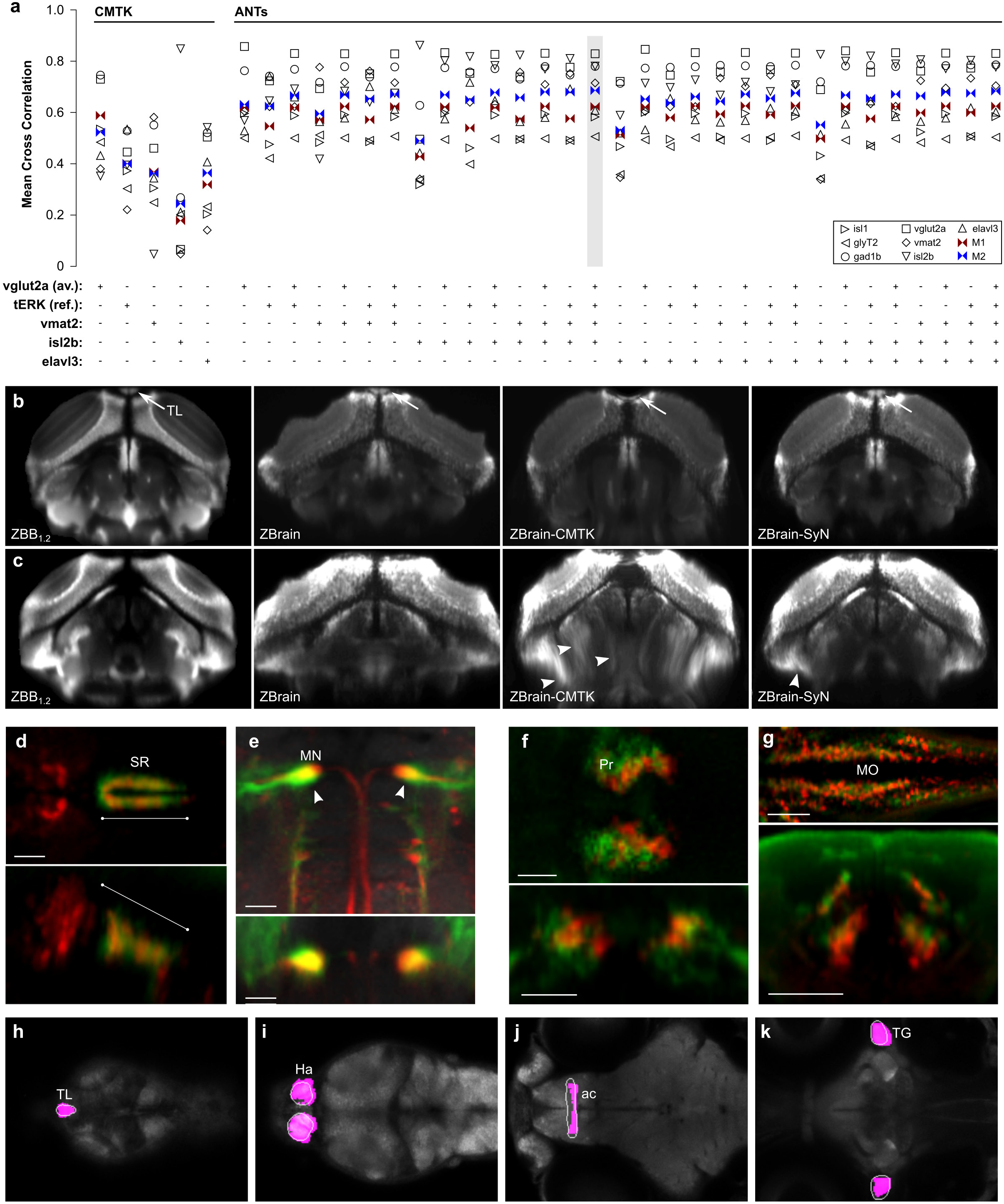
Transformation between Z-Brain and ZBB coordinate systems using multi-channel registration. (a) MCC for the expression patterns of *gad1b, glyT2, isl1, isl2b, tERK, vglut2a* and *vmat2* and the metrics M_1_ and M_2_, after registration of Z-Brain to ZBB_1.2_ using either CMTK or ANTs SyN with fixed-tissue registration parameters and the indicated combination of reference channels (*vglut2a, tERK_REF_, vmat2, isl2b and elavl3*). Note, similar results were obtained using the *tERK_AV_* instead of the *tERK_REF_* channel, but are omitted for clarity. The combination of reference channels selected for co-registration of Z-Brain and ZBB is highlighted. (b) Transverse view through the caudal optic tectum showing the *vglut2a* pattern in ZBB_1.2_, Z-Brain, Z-Brain after registration to ZBB with CMTK (ZBrain-CMTK), or with ANTs (ZBrain-SyN). The torus longitudinalis (TL) is well separated from tectal neurons in live scans, but less so in fixed tissue (arrows). The TL appears flattened after CMTK registration, but retains normal morphology after registration with ANTs SyN. (c) A comparison of transverse views as in (**b**), but slightly more caudal with contrast increased to highlight ventral distortion artifacts produced by registration (arrowheads). (**d-g**) Brain Browser views in the ZBB_1.2_ coordinate (**d,e**) or Z-Brain coordinate (**f,g**) space. Scale bars 25 μm except 50 μm in (**e**) (d) Horizontal (top) and sagittal (bottom) slices, comparing the *Pet1:GFP* expression pattern in the superior raphe in ZBB_1.2_ (red) and Z-Brain after transformation to the ZBB coordinate system (green). (e) Horizontal (top) and coronal (bottom) slices through the medulla oblongata, showing the expression of *y264Et* from ZBB_1.2_ (red) and *s1181Et* from ZBB-transformed Z-Brain (green), which both label the Mauthner cells (arrowhead). (f) Horizontal (top) and coronal (bottom) slice through the pretectum, comparing the expression of *DAT:GFP* from ZBB_1.2_ after transformation to Z-Brain coordinates (red) and anti-tyrosine hydroxylase staining in Z-Brain (green). (g) Horizontal (top) and coronal (bottom) slice through the medulla oblongata for *glyT2:GFP* from ZBB_1.2_ after transformation to Z-Brain (red) and the same transgenic line in Z-Brain (red). (**h-k**) Brain Browser horizontal slices showing manually segmented regions transformed from the Z-Brain coordinate system to ZBB_1.2_ (white outlines) compared to regions previously defined in ZBB obtained by thresholding expression patterns in transgenic lines (magenta). Regions are the torus longitudinalis (**h**), habenula (**i**), anterior commissure (**j**) and trigeminal ganglion (**k**).

## Results

### Optimization of ANTs based registration of live vglut2a:dsRed image scans

Brain registration in Z-Brain and ZBB used the B-spline elastic transformation in CMTK. Before attempting to co-align Z-Brain and ZBB, we tested an alternate algorithm for brain alignment, the diffeomorphic symmetric normalization (SyN) method in ANTs, because: (1) SyN has been shown to outperform B-spline transformations for deformable image registration in a variety of imaging modalities (Klein et al., 2009; Murphy et al., 2011). (2) ANTs permits registration using multiple reference channels, potentially allowing the use of complementary expression patterns as references for improved registration fidelity. (3) By calculating forward and reverse transformations simultaneously, SyN transformation matrices are intrinsically symmetric, ensuring that bridging registrations would be unbiased and that we could perform reciprocal transformations to register each dataset into the other’s coordinate system.

We first calibrated registration conditions by assessing alignment precision for a representative *vglut2a:DsRed* scan registered to the original *vglut2a:DsRed* reference brain in ZBB (*vglut2a*_ZBB_). Similar to CMTK we employed a three step registration within ANTs where rigid and affine steps were used to initialize a deformable registration using the SyN diffeomorphic transformation with cross correlation (CC) as the similarity metric. We tested a range of values for each of the SyN parameters as well as the radius of the region used for cross correlation.

While we previously used brain-wide normalized cross correlation (NCC) to evaluate registration (Marquart et al., 2015), correlation within local anatomical regions that contain discrete landmarks has been shown to be a more reliable criterion for quantitatively assessing registration precision (Rohlfing, 2012). Therefore, we identified a set of 12 landmarks within the *vglut2a* pattern, each within a 50 μm side cube. Landmarks were broadly distributed in the hope of representing diverse brain regions and minimizing the bias of any single structure. We measured the cross-correlation between the corresponding regions in *vglut2a*_ZBB_ and the registered image, then calculated the mean of the cross correlation between all regions (MCC; Fig. 1a). We also assessed the results visually to subjectively assess the severity of tissue distortion. Unsurprisingly, similar to our previous work with brain-wide NCC, images with the highest MCCs generally showed more conspicuous tissue distortion — thus although greater precision was achieved with increased deformation, we preferred results where cell shape and axon tract morphology were better preserved (Fig. 1b,c). Disregarding parameter combinations that resulted in overt distortion, we identified a set of values (Table 1, **live registration**) where cell morphology remained intact, but registration precision (MCC) was maximized. With these parameters, although the MCC for *vglut2a* improved only slightly from 0.79 using CMTK to 0.81 using SyN, cell morphology was noticeably better preserved, especially within ventral structures such as the hypothalamus (Fig. 1d).

We next tested whether these registration parameters also improved precision for the co-aligned transgenic lines. For ZBB, we co-scanned transgene and enhancer trap expression patterns with the *vglut2a:dsRed* transgene, allowing us to register each expression pattern to *vglut2a*_ZBB_. We first compared the overlap and morphology of the Mauthner cells from brain scans of three different individuals of transgenic line *J1229aGt* (Burgess et al., 2009). Overlap of Mauthner cell bodies was similar for CMTK and ANTs (Fig. 1e,f). However, in the CMTK registered images, the Mauthner axon was distorted in the caudal medulla, whereas axon morphology was normal with ANTs. Second, in our previous work, we assessed the precision of CMTK registration using line *y339Et*, by independently scanning two sets of three larvae, producing an average for each set, and visually comparing the result. With CMTK we had noted misalignment of approximately 1 cell diameter in the neuropil of the optic tectum (Fig. 1g). This was substantially improved with ANTs, where there was much closer alignment of the two averages (Fig. 1h). For quantification we calculated the cross correlation for 8 landmarks within the *y339Et* pattern, and found that the mean increased from 0.52 with CMTK to 0.63 with ANTs.

### Improved precision of ZBB after registration using ANTs

Based on the improved registration precision and reduced distortion of cell morphology achieved using SyN, we recompiled ZBB using ANTs for registration to create a more accurate atlas. We used ANTs to register the entire set of 354 brain scans that were part of ZBB, then as before, averaged multiple larvae to create a representation of each transgenic line, masked the average stacks to remove tissue outside the brain and re-imported the resulting images into our Brain Browser software. We will refer to this new recompilation of our atlas as ZBB_1.2_.

To determine whether ZBB_1.2_ was a quantitative improvement over ZBB, we calculated a cross-correlation score for each pattern in the browser. To avoid manually defining landmarks for each line, we instead computationally identified three regions inside each pattern with strong expression to act as landmarks. For each of these regions, we iteratively performed pair-wise cross-correlations between all individual brains from the same transgenic line, allowing us to calculate a mean cross-correlation (MCC) value for each line. We performed this procedure first for brains registered using CMTK, then for the same set of brains registered using ANTs, allowing us to compare MCCs for the two methods (Fig. 2a). Overall, the correlations increased slightly from ZBB to ZBB_1.2_ (0.32±0.02 to 0.34±0.02; paired t-test p=0.15). Although this was not statistically significant, it was instructive to examine instances with large changes in mean cross correlation. Line *y332Et* labels a small set of cells with a salt and pepper pattern in the right habenula. Here, cross correlation was greater after registration with CMTK (CMTK, 0.50; ANTs 0.39), due at least in part to greater distortion of cells resulting in increased overlap between individual fish despite the biological variability (Fig. 2b). In *y341Et*, distortion artifacts also appeared to account for the large increase in MCC obtained with ANTs (CMTK, 0.19; ANTs 0.58). Here, cells in the caudal hypothalamus had an elongated morphology after registration with CMTK, often stretching outside the boundaries of the nucleus. Consequently, this distortion reduced rather than increased the cross correlation score (Fig. 2c).

Additionally, we inspected regions of ZBB_1.2_ where we had noticed poor registration precision or pronounced cell distortion in the original ZBB. One such area was the dorsal thalamus, where cell morphology was noticeably perturbed after elastic registration with CMTK, with cell somas stretching across the midline (Fig. 2d). In ZBB_1.2_ cells retained a rounded morphology with distinct cell clusters on the left and right sides of the brain (Fig. 2e). Similarly, distortions in cell shape that were apparent in the caudal hypothalamus in ZBB, were absent in ZBB_1.2_ (Fig. 2f,g). In the caudolateral medulla, we previously obtained poor registration, with expression extending to regions outside the neural tube (Fig. 2h). In ZBB_1.2_, patterns had improved bilateral symmetry and were correctly confined to the neural tube (Fig. 2i). Finally, we noticed that the posterior commissure was poorly aligned between larvae leading to a defasciculated appearance in ZBB (Fig. 2j), whereas this tract had the correct tightly bundled appearance in ZBB_1.2_ (Fig. 2k).

Together, these observations confirm that ZBB_1.2_ is a more faithful representation of the transgenic lines. Not only is cell morphology better preserved, but metrics of global registration precision as measured by mean cross correlation are nevertheless improved from those of the original ZBB atlas.

### Optimization of ANTs registration parameters for fixed tissue

The Z-Brain atlas was derived by registering brain scans to a single brain that was fixed, permeabilized and immunostained for tERK expression. We therefore presumed that tERK would be a useful channel for bridging the two atlases, if we could first successfully register a tERK stained *vglut2a:DsRed* expressing brain to ZBB_1.2_. Therefore, we fixed and co-stained a transgenic *vglut2a:DsRed* larva for DsRed and tERK, and registered the tERK pattern to ZBB_1.2_ using the *vglut2a* pattern. We used the resulting image as our ZBB tERK reference brain (tERK_ZBB_).

In addition to the tERK reference brain, Z-Brain contains an average of 197 tERK stained larvae, which we thought might serve as a bridge between atlases. During studies on pERK-based activity mapping, we had generated a dataset of 167 tERK stained brains and sought to use these to create an average tERK representation by registering them to tERK_ZBB_. However, during this process, we noticed a high degree of variability between tERK stained brains, most notably in either poor labeling of ventral brain structures or in deformation of the optic tectum neuropil. Immunohistochemistry for tERK proved highly sensitive to staining parameters with the trypsin activity, permeabilization duration, and antigen retrieval having the strongest effects. This variability was most apparent in the optic tectum, where high trypsin activity tended to disrupt morphology and reduce the dimensions of the tectal neuropil (Fig. 3a,b). These local distortions were not corrected by deformable image registration: alignment to tERK_ZBB_ with the same parameters optimized for live *vglut2a* based registration, failed to correct the reduced tectal neuropil volume (Fig. 3c,d; asterisk) and often created an artifact where the neuropil zone failed to abut the underlying cellular layer labeled by *vglut2a* expression (Fig. 3c,d; arrowheads).

We therefore varied the registration parameters that were optimal for live *vglut2a* registration, to find settings that best rectified the variable tissue morphology following fixation and permeabilization. For tERK registration optimization, we used a set of 6 tERK stained brains including the Z-Brain tERK reference. We iteratively varied parameters for registration to tERK_ZBB_ and calculated the mean cross-correlation between each of the aligned tERK stains and tERK_ZBB_ (e.g., Fig. 3e,f). Again when visually inspected, we noted a trade-off between the quality of global alignment and local distortion artifacts, with the parameters which yielded the greatest increase in MCC often producing abnormally elongated cell profiles throughout the brain (Fig. 3g). However, visual inspection confirmed that parameters which increased MCC for fixed tissue greatly improved the morphology of the optic tectum neuropil (Fig. 3h,i). We therefore used ANTs with the fixed brain parameters (Table 1, **fixed registration**) to register 167 tERK stained brains to tERK_ZBB_, and generated an average tERK representation comparable to the Z-Brain tERK average (Fig. 3j,k).

### Inter-atlas registration using multi-channel diffeomorphic transformation

By chance, both Z-Brain and ZBB incorporated seven additional gene or transgene expression patterns that we judged of sufficient quality to act either as templates for bridging the datasets or to provide metrics for assessing the precision of the bridging registration (Table 2). For example, *vglut2a*_ZBB_ is a confocal scan of DsRed in a single larva from transgenic line *TgBAC(slc17a6b:loxP-DsRed-loxP-GFP)nns14*, whereas Z-Brain includes *Tg(slc17a6:EGFP)zf139*. In both cases, reporter expression is regulated by the same bacterial artificial chromosome (Bae et al., 2009; Satou et al., 2013). Crossing these two lines allowed us to scan DsRed and EGFP in the same larva and confirm that the patterns were largely congruous, potentially allowing us to use *vglut2a* expression to bridge the two atlases. Likewise, the expression patterns of *tERK, elavl3, isl2b, vmat2* in Z-Brain and ZBB appeared sufficiently similar to provide templates for atlas co-registration.

**Table 2.**
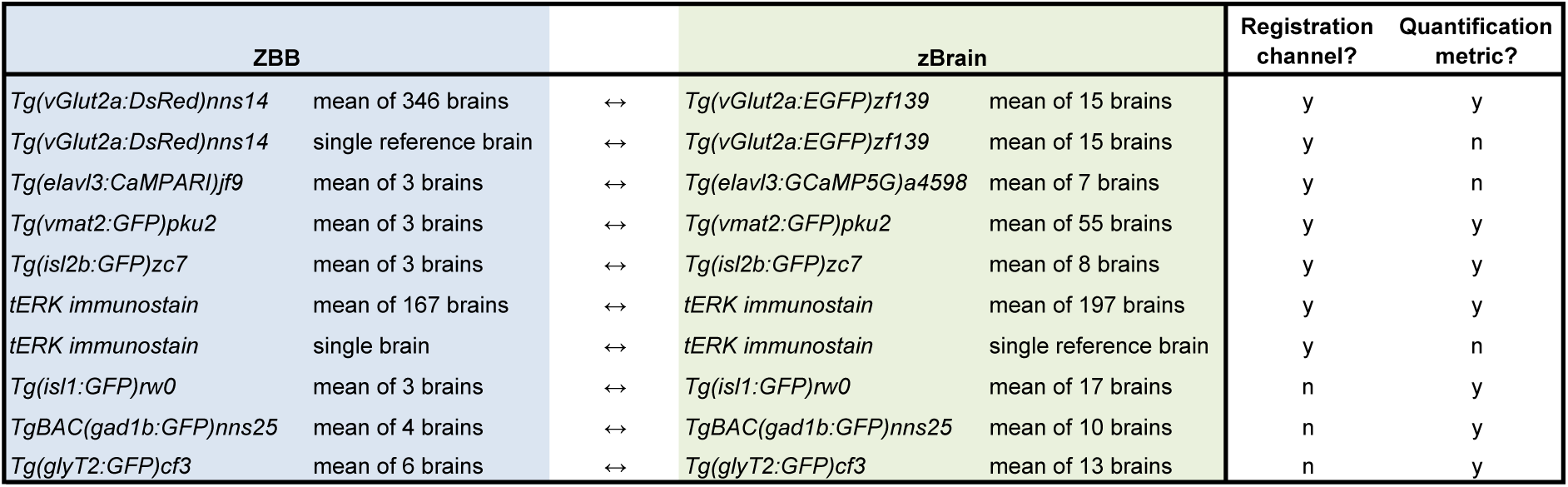
Brain images in ZBB and Z-Brain that were used as templates for registration and/or for measurement of registration precision.

We used seven expression patterns to evaluate registration precision: *vglut2a*, *isl2b, vmat2, elavl3, isl1, gad1b* and *glyT2*. For each pattern we identified a set of 5–18 landmarks that were widely distributed to represent diverse brain regions. For each landmark, we measured the cross-correlation between the corresponding volumes in ZBB and Z-Brain. We then calculated the mean of all cross correlation (MCC) values for landmarks associated with a given expression pattern. We used two measures of registration precision. The first metric (M_1_) was the mean of the MCCs for *isl1, gad1b* and *glyT2* expression patterns in ZBB and in Z-Brain after registration to ZBB. These three expression patterns do not provide sufficient coverage across all brain regions to use for registration, but served as independent channels to estimate registration precision. However, as these patterns are relatively sparse they do not comprehensively assess precision across all brain regions. To provide a global measure of precision, we therefore also used a second metric (M_2_) that was the mean of all seven MCCs: those in M_1_ plus four of the patterns used as references for registration - *vglut2a, tERK, isl2b and vmat2*. Although M_2_ uses expression patterns that together provide good coverage for the entire brain, we expected that the four patterns that were also used to guide the deformable registration, would artificially inflate the MCC.

We first used CMTK to register Z-Brain to ZBB_1.2_. Maximal M_1_ and M_2_ scores were obtained using the average *vglut2a* pattern as the reference (Fig. 4a). We therefore registered all images in Z-Brain to ZBB using the *vglut2a* average in each dataset as the reference channel. We observed severe tissue distortions in several brain regions, with noticeable flattening of the torus longitudinalis and tissue distortions, particularly in ventral brain regions (Fig. 4b,c; ZBrain-CMTK).

Next, for comparison, we used the ANTs SyN algorithm to register the atlases. Ideally, patterns for registration should include information throughout the brain. Because ANTs can use multiple concurrent reference channels to derive an optimal transformation matrix, we speculated that the best possible transformation would be achieved by a combination of channels with complementary information. We therefore produced an inter-atlas transformation matrix using every combination of the *elavl3, isl2b, vglut2a, vmat2, tERK*_*REF*_ (tERK single brain) and *tERK*_*AV*_ (tERK average brain) patterns as references. As Z-Brain used fixed samples, we used the registration parameters previously optimized for the greater variability present in fixed tissue. Multi-channel registration significantly improved M_1_ and M_2_ values compared to any single channel alone and to transformations obtained using CMTK. The registration obtained with *vglut2a, tERK*_*REF*_, *vmat2* and *isl2b* gave the highest M_2_ value and an M_1_ score within 1% of the highest scoring combination (Fig. 4a). Moreover, the overt tissue distortions noted after elastic registration with CMTK were far less salient using these parameters (Fig. 4b,c; ZBrain-SyN). We therefore applied the transformation matrix obtained with this set of channels to the database of gene expression patterns in Z-Brain to align them to ZBB_1.2_.

The precision of the inter-atlas registration is apparent when comparing the location of cells that are present in both datasets, such as those labeled by *Pet1:GFP*. The Z-Brain transformed pattern closely matches the transgene expression pattern in ZBB_1.2_ within the superior raphe (Fig. 4d — note however that unexpectedly, the line in ZBB_1.2_ also labels a set of more rostral cells not apparent in Z-Brain). Both atlases also include lines labeling the Mauthner cells. After registration, Mauthner cells in the atlases substantially overlapped, although they were several microns more medially positioned in ZBB_1.2_ (Fig. 4e). Similarly, we used the inverse of the transformation generated by SyN to register ZBB_1.2_ to the Z-Brain coordinate system. As expected, expression in the *DAT:GFP* line in ZBB_1.2_ overlapped well with the tyrosine hydroxylase stain from Z-Brain in the pretectum (Fig. 4f), although again, the ZBB_1.2_ pattern was slightly more medial than in Z-Brain. More caudally, the *glyT2:GFP* transgenic line labels glycinergic neurons in longitudinal columns in the medulla oblongata (Kinkhabwala et al., 2011). These columns were closely aligned after ZBB_1.2_ was registered to Z-Brain (Fig. 4g).

Z-Brain includes 294 masks that represent anatomically defined brain regions or discrete clusters of cells present in transgenic lines. We selected 113 of these masks that delineate neuroanatomical regions and transformed them into the ZBB_1.2_ coordinate system. We had previously defined a small number of our own anatomical masks by thresholding clusters of neuronal cell bodies located in well-defined brain regions. In contrast, the Z-Brain masks are more comprehensive, have smoother boundaries and include both the cell bodies and neuropil for a given region (Fig. 4h-k). We therefore imported the Z-Brain masks into ZBB_1.2_, replacing most of our existing masks. We also modified the *Brain Browser* software to automatically report the neuroanatomical identity of a selected pixel, or to display the boundaries of the region encompassing a selected point. The updated software and rebuilt database in ZBB_1.2_ can be downloaded from our website (https://science.nichd.nih.gov/confluence/display/burgess/Brain+Browser).

## Conclusions

Digitized data-derived brain atlases provide an opportunity to continuously integrate new information and iteratively improve data accuracy within a common spatial framework. Thus, as methods evolve and technology improves, insights can be easily added to existing data to provide an increasingly rich view of brain structure and function. Because the entire larval zebrafish brain can be rapidly imaged at cellular resolution, it is possible to envisage an atlas that combines detailed information on cell type (including gene expression and morphology), connectivity and activity under a variety of different physiological conditions. At present, biological variability presents an obstacle, as brain regions contain multiple intermingled cell types that are not positioned in precisely the same manner between larvae. To circumvent this in the existing zebrafish brain atlases, multiple individuals of a given line are sampled and averaged to generate a representative expression pattern. Current atlases are thus essentially heat maps of gene expression or activity. Despite this spatial ambiguity, aggregating information from different sources into the same spatial framework still provides valuable indicators of cell type, gene co-expression, and neural activity under defined conditions.

Ideally different atlas projects might use the same reference brain, however in practice the choice of a reference is dictated by study-specific requirements. For example, despite the deformations introduced by fixation and permeabilization, a fixed brain is essential for activity mapping using pERK immunohistochemistry. In contrast, we were able to take advantage of the optical transparency of larvae to rapidly scan and register several hundred individuals representing more than 100 different transgenic lines. For our purposes, the *TgBAC(slc17a6b:loxP-DsRed-loxP-GFP)nns14* line was ideal, because through Cre injection, we generated a *vglut2a:GFP* line with an almost identical pattern, allowing us to co-register lines with either GFP or RFP fluorescence. However, we have also used pan-neuronal Cerulean or mCardinal as a reference channel when the green and red channels both contained useful information on transgene expression. Our work now demonstrates that it is feasible to contribute to community efforts at building an integrated map of brain structure, expression and activity, while allowing reference image selection to be guided by technical considerations.

One caveat to this conclusion is that deformable image registration can easily introduce artifacts into cell morphology if parameters are not carefully monitored and constrained. Indeed, a special challenge for brain registration in zebrafish is preserving the local morphology of neuronal cell bodies and axons, while permitting sufficient deformation to correct for biological differences and changes in brain structure arising from tissue fixation and permeabilization. Thus, while B-spline registration with CMTK produced acceptable inter-atlas alignment, it also introduced noticeable distortions into local brain structure that affected neuronal cell morphology. Such artifacts were particularly severe in ventral brain regions such as the caudal hypothalamus, and may therefore be due to differences in ventral signal intensity between the datasets. In ZBB, in order to compensate for the increase in light diffraction with tissue depth, we systematically increased laser intensity with confocal scan progression (z-compensation). As a result, the Z-Brain and ZBB datasets are comparable in dorsal brain regions, but there is a noticeable discrepancy ventrally which may account for the loss of registration fidelity. Alternatively, although z-compensation partially corrects for reduced fluorescent intensity, there is a noticeable drop-off in image resolution in ventral regions; the resulting loss of information may lead to lower quality registration. Registration algorithms that allow parameters to vary by depth may ameliorate these physical imaging constraints.

Nevertheless, the symmetrical diffeomorphic transformation in ANTs provides a satisfactory solution to these problems. For live tissue, we found parameters that allowed the ANTs SyN transform to achieve similar or better registration precision than previously achieved using CMTK, while minimizing overt distortions in tissue structure and neuronal cell morphology. In our hands, permeabilization of fixed tissue tended to produce variable changes in neuropil structure which was most salient in the optic tectum.

Specifically, neuropil volume was diminished when fresh aliquots of trypsin were used for extended durations. These artifacts can be minimized by stringent oversight of reagent viridity. However, by calibrating SyN parameters to permit larger deformations, we were able to accommodate the variability introduced by tissue processing.

The main limitations for use of the SyN registration algorithm in ANTs are the large memory demands (73 GB for a single channel registration) and long computational times (3-5 hours for a single channel using 24 cores) required for registration of images with a resolution sufficient for the brain-wide visualization of neuronal morphology (e.g., 1000 × 600 × 350 pixels). For multi-channel registrations, memory demands and computation time were even greater: 106 GB for 6 channels taking over 16 hours on 24 cores. However, our present ANTs SyN parameters likely can be further optimized to reduce these demands. For instance, our parameters currently include 10 iterations of transformation matrix optimization at full image resolution. In our experience, these full resolution registration cycles do not significantly improve cross correlation scores, but greatly increase computation time. Thus, computation time may be reduced by adjusting registration resolution as well as other parameters without adversely affecting registration quality. Although computational resources did not present a bottleneck for registering a small number of samples, this increase in the demands of an individual registration made it difficult to optimize registration parameters as extensively as we had done previously with CMTK (Marquart et al., 2015). For example, during our initial effort to optimize registration parameters for live *vglut2a* expression, we used a single representative example rather than assessing parameters for a set of several independent scans. By reducing computation time, we would be able to explore more comprehensively the parameter space available with SyN, and evaluate alternative diffeomorphic transforms available with ANTs that may provide still better registration fidelity.

An obstacle to systematically calibrating registration parameters was finding a suitable metric to quantitatively evaluate registration precision. This is a recognized problem, and it is not clear that a general solution exists (Rohlfing, 2012). We used cross-correlation within image neighborhoods that included relatively high contrast internal image boundaries. However in registering live *vglut2a:DsRed* image stacks, we found that the highest scoring transformations achieved accurate global brain alignment at the expense of biologically plausible cell morphology. Therefore, it was essential to visually compare the output of every transformation and make subjective judgments about registration quality. This was difficult, because distortions, when present, tended to be variable in different parts of the image, thus requiring the entire image stack produced by each transformation to be scrutinized to select optimal parameter settings.

Nevertheless, this study demonstrates that the Ants' diffeomorphic symmetric normalization algorithm improves upon elastic registration for precise registration of whole brain images in larval zebrafish and is markedly better at preserving neuronal cell morphology. By systematically testing SyN registration parameters for registering images acquired using live scans, we improved the ZBB atlas. Then, after calibrating registration parameters for fixed tissue and using multi-channel optimization, we were able to align the Z-Brain atlas into the ZBB coordinate space, and vice-versa. We believe that integrating the information present in each of these atlases produces a richer framework for future studies of structural and functional relationships within the nervous system. Large digital datasets such as those present in brain atlases can be used for many types of bioinformatic analysis. Z-Brain and ZBB already include software that can be used to explore the larval zebrafish brain, and we hope that by integrating these datasets into a single coordinate system, we will help to stimulate the development of additional computational tools and methods for querying this information.

## Acknowledgements

This work was supported by the Intramural Research Program of the *Eunice Kennedy Shriver* National Institute for Child Health and Human Development (NICHD) and utilized the high-performance computational capabilities of the Biowulf Linux cluster at the National Institutes of Health, Bethesda, MD (https://hpc.nih.gov). We thank Owen Randlett for valuable discussion and help checking the correspondence of *vglut2a* expression patterns. We are grateful to Sinisa Pajevic (NIH/CIT) for advice on computational procedures and to M. Okan Irfanglu and Neda Sadeghi (NICHD) for guidance in optimizing ANTs parameters.

